# Evaluating flash freezing for preservation of rat abdominal aorta for delayed biomechanical characterization

**DOI:** 10.1101/2023.10.16.562465

**Authors:** Koen W.F. van der Laan, Koen D. Reesink, Sara Lambrichts, Nicole J.J.E. Bitsch, Laura van der Taelen, Sébastien Foulquier, Tammo Delhaas, Bart Spronck, Alessandro Giudici

## Abstract

Most studies investigating arterial stiffening use animal rather than human arteries. This is because human tissue becomes available in small amounts and at irregular times, which complicates planning of experimental work. Suitable tissue preservation methods for delayed biomechanical testing prevents the need for testing fresh tissue and alleviates some of the logistical challenges of human *ex vivo* studies. Therefore, the present study aimed to investigate whether the existing method of flash freezing and subsequent cryostorage provides is suitable for delaying the characterization of arterial biomechanics. Fresh and flash frozen abdominal aortas (*n*=16 and 14, respectively) were quasi-statically and dynamically tested using a biaxial testing set-up with dynamic pressurization capabilities. The acquired biomechanical data was modeled using a constituent-based quasi-linear viscoelastic modeling framework, deriving directional stiffness parameters, individual constituent biomechanical contributions, and viscoelastic stiffening under dynamic pressurization conditions. Flash freezing reduced arterial wall thickness, increased circumferential stiffness, as well as reduced viscoelastic stiffening at higher pressures. These findings reflected those in the modeled contribution of collagen to arterial biomechanics, showing increased collagen load bearing at higher pressures. However, despite the above mentioned detectable changes, flash freezing did not alter the mechanical relation between elastin and collagen, maintaining a non-linear response to pressurization and stretch. Flash freezing may thus be suitable for studies requiring delayed characterization of passive arterial biomechanics, assuming care is taken to ascert that the impact of flash freezing on study groups can be approached as a systematic error.

## 1 INTRODUCTION

Arterial stiffening is an important risk factor for cardiovascular disease, a leading cause of death globally, and the subject of extensive research [1-5]. Pulse wave velocity (PWV) is commonly used for *in vivo* characterization of arterial stiffening in human and animal studies, but is insufficient for underpinning the underlying root causes for changes in arterial elastic properties [6]. The complex nature of the arterial wall tissue requires performing *ex vivo* experiments to disentangle the interactions between its different constituents (e.g., elastin, collagen and vascular smooth muscle cells (VSMCs)) to explore the potential causes of arterial stiffening. It is important to note that, because biological tissues and their properties decay over time, these experiments are typically performed within hours after tissue excision. However, the use of fresh tissue poses logistical challenges. First, proximity between the tissue source and the testing facility is necessary to avoid excessive delays due to transportation. Second, sacrifice rates should not exceed the processing capacity of the testing facility. Third, lab equipment and personnel must be available when tissue becomes available. Human *ex* vivo studies exacerbate some of these challenges, as human arterial tissue is scarce and comes available in small amounts and at irregular times. Effective tissue preservation methods that maintain tissue biomechanics for prolonged periods would help reduce these logistical challenges.

Tissue sample freezing is a common preservation method to enable long term storage of biological tissues. However, care should be taken when selecting freezing methods, as there is the potential to permanently damage or alter tissue properties during the preservation process [7]. Ice crystal formation is the primary cause of damage during freezing, by puncturing surrounding structures, dehydrating cells, and generating osmotic shock and potential cell rupture. Indeed, previous studies have shown that freezing arterial tissue results in loss of VSMC contractility and viability, as well as damage to extracellular matrix (ECM) components, such as collagen and elastin fibers [8-10]. Tissue damage could potentially be reduced by limiting ice crystal formation, either by rapidly cooling or, in contrast, by decreasing the cooling rate and adding cryopreservatives, which minimize ice crystal formation [11]. Several studies have demonstrated that arterial cell viability and contractility can be partially preserved with slow cooling rates and added cryopreservatives [12-14]. Furthermore, histological evidence shows that these techniques have little or negligible impact on the ECM [15]. Of note, during thawing care should also be taken to slowly thaw frozen arterial tissue, because rapid thawing might cause fractures [13, 14, 16].

Few studies investigated the impact of cryopreservation on arterial biomechanics, whilst reporting varying results [15, 17-19]. Planar biaxial testing has shown increased stiffness at high deformations for cryopreserved arterial tissue compared to fresh tissue [17, 19]. Wire myography revealed overall stiffening of arterial tissue with cryopreservation [15]. In contrast, pressure myography showed that cryopreservation had a softening effect on arterial tissue [18]. Direct comparisons between existing studies are hampered due to differences in cryopreservation methods, which may differentially affect arterial tissue structure and mechanics, as well as by the use of different mechanical testing methods. In addition, while the impact of cryopreservation on quasi-static arterial elastic properties has been investigated, the impact on dynamic arterial elastic properties, which are more relevant to in vivo pressure pulsatility, remains yet to be explored.

The aim of this study was to determine the impact of flash freezing and subsequent cryostorage at -80°C on static as well as dynamic arterial elastic properties, thereby evaluating the suitability of this preservation method for delayed testing of arterial biomechanical properties. We conducted our study by biomechanically characterizing rat abdominal aortic segments under *in vivo*-like conditions. Arterial wall constituent contributions to the whole-wall biomechanics were disentangled using a constituent- based modelling approach [20]. Arterial wall collagen microstructure was imaged using two photon laser scanning microscopy (TPLSM), to be able to correlate changes in microstructure to changes in biomechanical properties.

## 2 METHODS

### 2.1 Sample preparation

Abdominal aortas were obtained from lean (*n*=10) and obese (*n*=10) ZSF1 rats at 22 to 23 weeks of age (Charles River Inc., Wilmington, MA, United States). All experimental protocols and methods involving animals within this study were conducted in accordance with institutional guidelines and approved by the Maastricht University Animal Ethics Committee (AVD10700202010326). Animals were terminated by an overdose of pentobarbital (100 mg/kg, i.p.) under isoflurane anesthesia. Abdominal aortas (AAs) were excised, placed in calcium- and magnesium-deficient Hank’s balanced salt solution (HBSS) buffer and cleaned from connective tissue and fat. Samples were cut into a proximal and a distal tubular segment, typically divided at the inferior mesenteric artery (IMA). Half of the lean and obese proximal and distal AA segments were allocated to the fresh and frozen groups, respectively, creating eight subgroups with equal sample sizes (*n*=5). Paper tissue was used to remove excess buffer from the samples undergoing cryopreservation. These samples were subsequently placed inside a dry Eppendorf tubes and closed off. The Eppendorf tubes were then submerged in liquid nitrogen, to flash freeze the samples inside. Frozen samples were stored for four weeks at -80 C°. Fresh samples were immediately stored in fresh calcium- and magnesium-deficient HBSS buffer at 4 C° and biomechanically tested within the next five hours.

### 2.2 Biomechanical characterization

#### 2.2.1 Biomechanical testing set-up

The biaxial biomechanical testing set-up used in this study (**Figure 1**) is a custom-built pressure myograph based on a previously reported set-up for investigating tissue biomechanics of large vessels of small animal models [21]. Compared to the previously described setup, the new setup: (1) tracks diameter using a high speed camera (USB 3 uEye CP Rev. 2, IDS Imaging Development Systems GmbH, Obersulm, Germany) instead of a high-frequency ultrasound transducer, allowing for diameter tracking during both inflation and stretching experiments, (2) facilitates dynamic sample inflation with a custom pressure oscillator, allowing for customized pressure pulses with a frequency range of 0.1 to 20 Hz, and (3) has both proximal and distal cannulation pipettes mounted on motorized stages, allowing for controlled axial stretching of the samples (i.e., along the vessel main axis) whilst keeping the center of the sample within the camera’s field of view. Using this set-up, samples were cannulated on glass pipettes in the heated organ bath (37°C), then inflated and stretched to investigate their biaxial biomechanics.

**Figure 1:**
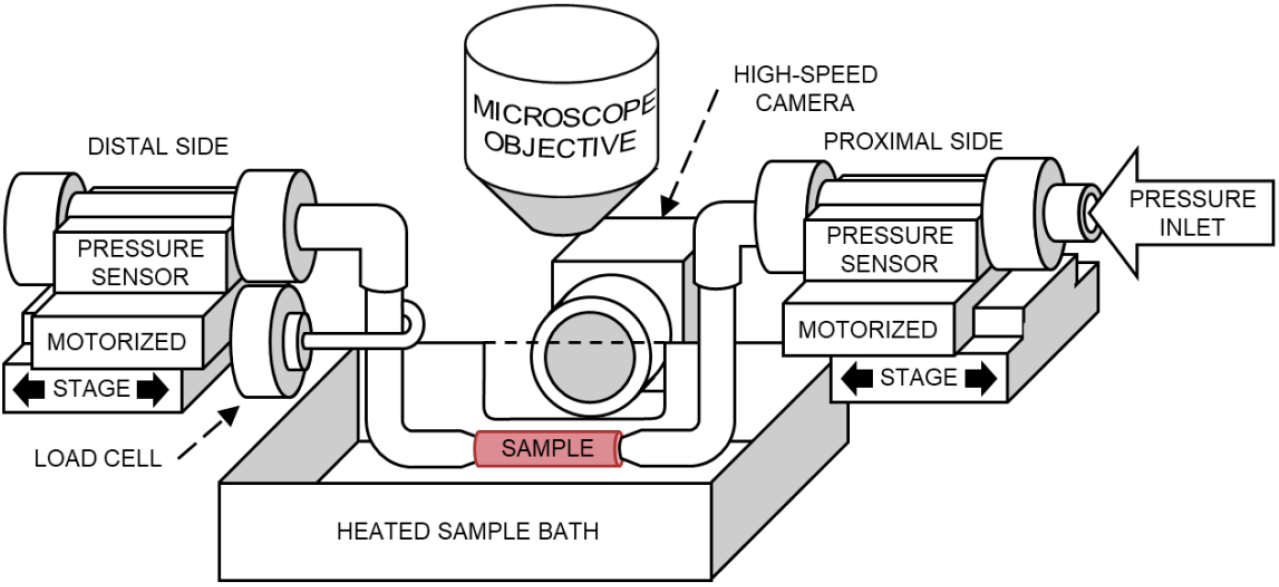
Biaxial biomechanical testing set-up. Samples are mounted on glass pipettes held inside a heated sample bath. A transparent glass slide on one side of the bath allows for tracking the outer diameter via a (high- speed) camera during pressurization and axial stretching.

#### 2.2.2 Biomechanical testing protocol

After cryogenic storage, sample containers were allowed to reach room temperature in the open air. Samples were then submerged in HBSS; samples stored in HBSS at 4 °C were placed in new HBSS buffer. Subsequently, side branches were sutured closed. The organ bath, buffer reservoir and tubes in the set-up were then filled with HBSS, taking care to remove any air bubbles in the system of tubing. 10 μM sodium nitroprusside (SNP) was added to the organ bath to minimize VSMC contraction during mechanical testing. After the organ bath reached 37 °C, samples were cannulated onto the glass pipettes and brought to an unloaded axial length at which samples were visually neither stretched nor buckling.

First, vessels were axially stretched to their *in vivo*-like length, at which the minimum change in axial force was detected by the load cell while luminal pressure oscillated between 10 and 140 mmHg. Next, for circumferential preconditioning, samples were axially stretched to 105% of their *in vivo*-like axial stretch and then slowly pressurized from 10 to 180 mmHg and back for four times. Finally, for longitudinal preconditioning, samples were pressurized to 100 mmHg and slowly stretched four times from zero to the maximum axial force noted during circumferential preconditioning and back, using the motorized stages. Samples were then depressurized and returned to an unloaded state, judged visually. The unloaded length of the sample (i.e., the length between the cannulation points) was then recorded and its *in vivo*-like axial stretch was redetermined.

The biomechanical characterization protocol comprised three series of experiments. First, samples were pressurized from 10 to 180 and back to 10 mmHg at a rate of 3 mmHg/s, with a pause of two seconds at maximum pressure. Measurements were repeated twice at 105%, 95% and 100% of the *in vivo*-like axial stretch. Second, samples were pressurized dynamically, utilizing the pressure oscillator to generate sinusoidal pressure waves with an amplitude of 20 mmHg and frequencies of 0.625, 1.25, 2.5, 5, 10, and 20 Hz. For each frequency, harmonic pressurization experiments were performed at three mean pressures of 60, 100, and 140 mmHg. Third, quasi-static force-length experiments were performed by stretching the sample from zero to maximum axial load, as recorded during the quasi-static pressurization experiments, and back to zero load. Samples were stretched with a rate of 0.0187 s^-1^ with respect to the unloaded length. Stretching experiments were repeated twice at each of the following pressures: 10, 60, 100, 140 and 180 mmHg.

#### 2.2.3 Two-photon laser scanning microscopy imaging

After biomechanical characterization, the setup was placed under a Radiance2100 two-photon laser scanning microscope (TPLSM) (Bio-Rad, Hercules, CA, United States). Samples were stretched to their *in vivo*-like length and pressurized to 100 mmHg. Using a 60x CFI APO NIR Objective (NA 1.0, WD 2.8 mm) (Nikon, Minato City, Tokyo, Japan), second harmonic generation (SHG) image stacks of the sample were recorded using a Tsunami tunable pulsed femtosecond laser (Spectra-Physics, Santa Clara, California, United States) set to an 810 nm wavelength. SHG light originating from collagen fibers was isolated using a 400-410 nm high quality fluorescent filter and captured with a photon multiplier tube. One Image stack was recorded per sample with a field of view of 205 × 205 μm^2^ and an image resolution of 1024 x 1024 pixels^2^, resulting in a pixel size of 0.2 x 0.2 μm^2^. For each stack, images were recorded with 0.45 μm vertical steps between them until all contrast was lost.

#### 2.2.4 Cross-section imaging

After TPLSM imaging, the samples were depressurized and returned to their unloaded length. Three thin cross-sectional rings were cut from the sample. The three rings and a small ruler were placed in a Petri dish filled with HBSS. Cross-sectional view images of the rings were taken using a USB camera (Dino-lite, Almere, The Netherlands) mounted in a stereo microscope and wall thickness was determined using a custom MATLAB (R2022a, MathWorks, Natick, MA, United States) script, as reported previously [21].

#### 2.2.5 Model-derived biomechanical characterization

We used our recently developed constituent-based quasi-linear viscoelastic (cbQLV) modeling approach to quantify the arterial wall biomechanical behavior [20]. Briefly, rat aorta is assumed to be a thin-walled cylinder, the passive viscoelastic mechanical behavior of which is determined by the superimposed contributions of elastin and collagen. The model utilizes a homeostatic *in vivo* reference configuration, defined as the experimentally determined vessel configuration at the *in vivo*-like axial length while pressurized at 100 mmHg [22]. The model parameters were fitted to the acquired quasi-static and dynamic experimental data and the fitted model was used to simulate the vessel’s response to pressurization while being held at the *in vivo*-like axial stretch. For these simulations, the vessel-specific *in vivo*-like axial length was re-estimated as the crossover point of the axial force-axial length relationships measured during the quasi-static force-length experiments at 60, 100 and 140 mmHg of luminal pressure. Relevant geometrical features and biomechanical metrics were determined at 60, 100 and 140 mmHg. In addition, parameters influenced by dynamic pressurization, i.e., dynamic to quasi-static circumferential stiffness ratio and PWV, were determined at a pressure oscillation frequency of 10 Hz, physiologically the most relevant considering the resting heart rate in a conscious unrestrained mouse is around 600 BPM, at the pressure ranges 40–80, 80–120, and 120–160 mmHg [23].

#### 2.2.6 Microscopy image stack analysis

We used a custom MATLAB program to quantify the thickness of the collagen layer and the collagen volume fraction in the SHG image stacks. First, a gaussian filter with a 5×5 kernel was applied to the SHG channel image stack to increase the signal-to-noise ratio. Image intensity values were then adjusted so that 0 to 99.9% of voxels starting with the lowest intensity values filled the entire intensity range. A binary collagen mask was constructed from the image stack using voxels that exceeded a background threshold of 11.8% of the image intensity range. The sample’s centerline was determined by fitting a cylinder to this structure using a previously reported cylinder fitting method [24]. Radial positions for each voxel in this structure were determined with respect to the sample’s centerline while radial boundaries of the collagen layer were set to exclude the 1% smallest and largest voxel radial positions, as shown in **Figure 2**.

**Figure 2:**
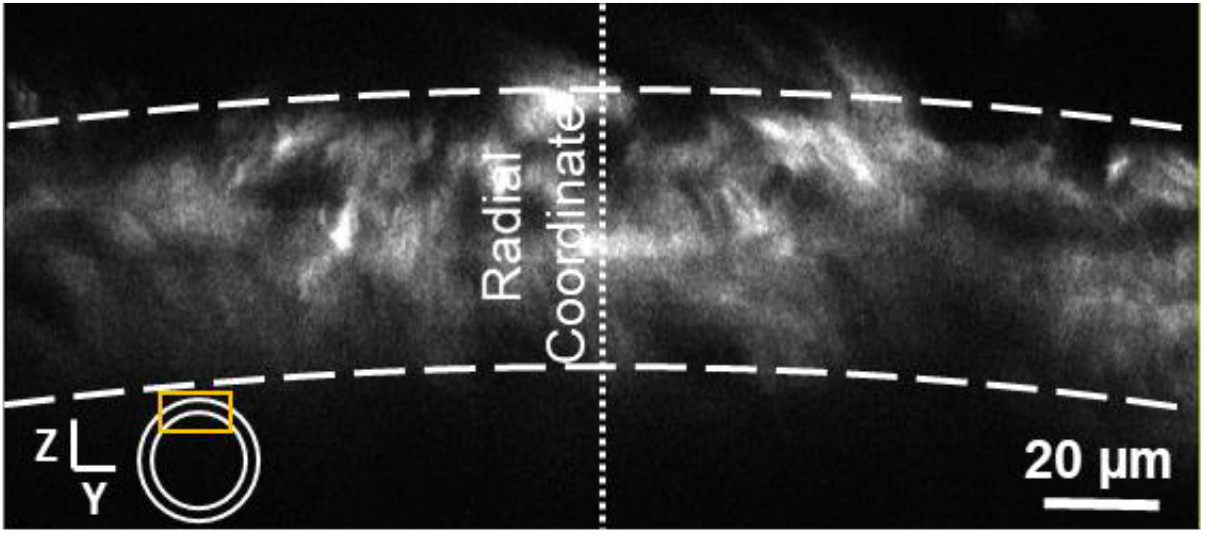
YZ slice from second harmonic generation image stack, represented by the yellow rectangle, illustrating the cross section of the collagen layer in the arterial wall. The dotted line represents the radial coordinate with respect to the vessel center line, dashed lines represent inner and outer radial boundaries of the collagen layer.

The volume and volume fraction of collagen inside image stacks was determined by the volume of voxels included in the collagen structure and the ratio between this volume and the total volume of the vessel wall inside the field of view of an image stack. The volume of the vessel wall inside the image stack’s field of view is determined from the inner and outer radius of the vessel at experimental *in vivo*- like stretch while pressurized to 100 mmHg, assuming the vessel lies flat in the field of view, oriented along the x-axis and centered in the y-axis with respect to the scanning mirrors of the microscope.

### 2.3 Statistical analysis

Three-way analysis of variance (ANOVA) was used to determine whether there were significant differences between fresh and frozen samples, because of the three independent variables along which samples could be divided into subgroups (i.e., fresh/frozen, lean/obese, and proximal/distal segment). A type III three-way ANOVA analysis was used to give equal weight to subgroups within fresh and frozen groups. Such an approach is more robust against potential sample dropout, as equal weighted subgroups prevents differences between subgroup sample sizes from influencing differences between fresh and frozen groups, instead resulting in an overall loss of statistical power [25]. Differences between fresh and frozen samples were considered statistically significant if *p*<0.05 and in the absence of significant interactions terms between independent variables that included the storage condition. In accordance with the statistical analysis method, the results are given as equal weighted means for fresh and frozen groups, as well as standard deviations determined with respect to the equal weighted means.

## 3 RESULTS

### 3.1 Sample dropout

During testing, failure to maintain luminal pressure for some samples due to leakage caused sample dropout and resulted in the group sizes shown in **Table 1**.

**Table 1:** number of successfully tested samples for each subgroup.

|  | Lean |  | Obese |  | Total |
| --- | --- | --- | --- | --- | --- |
|  | Proximal | Distal | Proximal | Distal |  |
| Fresh | 5 | 4 | 5 | 2 | 16 |
| Frozen | 4 | 5 | 2 | 3 | 14 |

Prox. AA: proximal abdominal aorta (i.e., above the inferior mesenteric artery); dist. AA: distal abdominal aorta (i.e., below the inferior mesenteric artery).

### 3.2 Effect of flash freezing on static response to inflation

The modeling approach captured the samples’ biomechanical behavior well, producing average coefficients of determination (*R*^2^) of 0.989 ± 0.0072 (mean standard ± deviation) and 0.995 ± 0.002 when modeling the static and dynamic biomechanical behavior, respectively, with no significant differences between fresh and frozen groups. The model parameters of the cbQLV model of all tested arteries are reported in Supplementary material, **Table S1**.

All statistically tested model parameters, along each independent variable (i.e., fresh/frozen, lean/obese and proximal/distal segment), are reported in the Supplementary material **Tables S2-4**. While fresh and frozen samples displayed similar responses to axial stretching, **Figure 3** shows different static responses to inflation for fresh and frozen samples at *in vivo*-like axial length. Frozen samples displayed a steeper pressure–diameter curve compared to fresh samples, suggesting cryopreservation stiffened the arterial wall. Fresh and frozen pressure–diameter mean curves crossed over each other at ∼90 mmHg, as fresh samples displayed smaller and larger diameters at low and high pressures, respectively.

**Figure 3:**
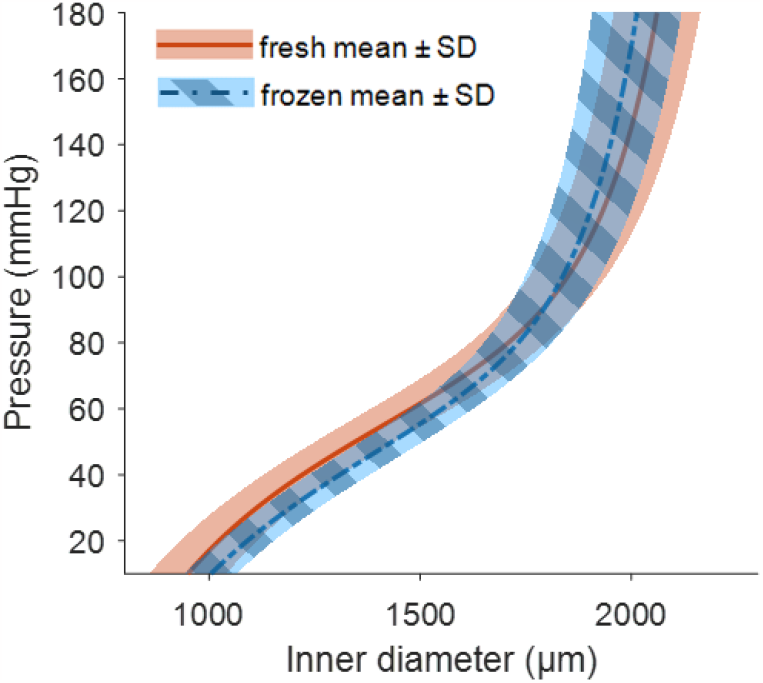
Model-derived static pressure-diameter relationship of fresh and flash frozen groups show a crossover, with frozen samples displaying above and below 92 mmHg reduced and increased diameters, respectively. Lines depict the mean static pressure-diameter relations (averaged in the diameter direction) at in vivo-like stretch, with equal weight given to subgroups within fresh and frozen groups. Shaded areas indicate standard deviation (SD).

**Figure 4B** shows that flash freezing of aortic tissue significantly reduced the unloaded wall thickness by about 15%, but preserved unloaded outer diameters (**Figure 4A**). When axially stretched and pressurized to *in vivo*-like stretch and 100 mmHg, respectively, wall thickness appeared reduced in flash frozen samples, though the difference was not significant (**Figure 4C**). Circumferential stretch of frozen samples, with respect to the loaded configuration, was significantly less (**Figure 4D**). Small deviations from the reference stretch at 100 mmHg for both groups arise from differences in experimentally determined and modeled *in vivo*-like stretches (i.e., simulations were run imposing the *in vivo*-like axial elongation determined *a posteriori* from the acquired data, while the reference configuration was imposed based on the *in vivo*-like axial elongation determined experimentally). Together with a reduced circumferential stretch, frozen samples displayed significant increases in circumferential stiffness, in collagen circumferential load bearing at higher pressures, and in PWV calculated from static data (**Figure 4E-G**).

**Figure 4:**
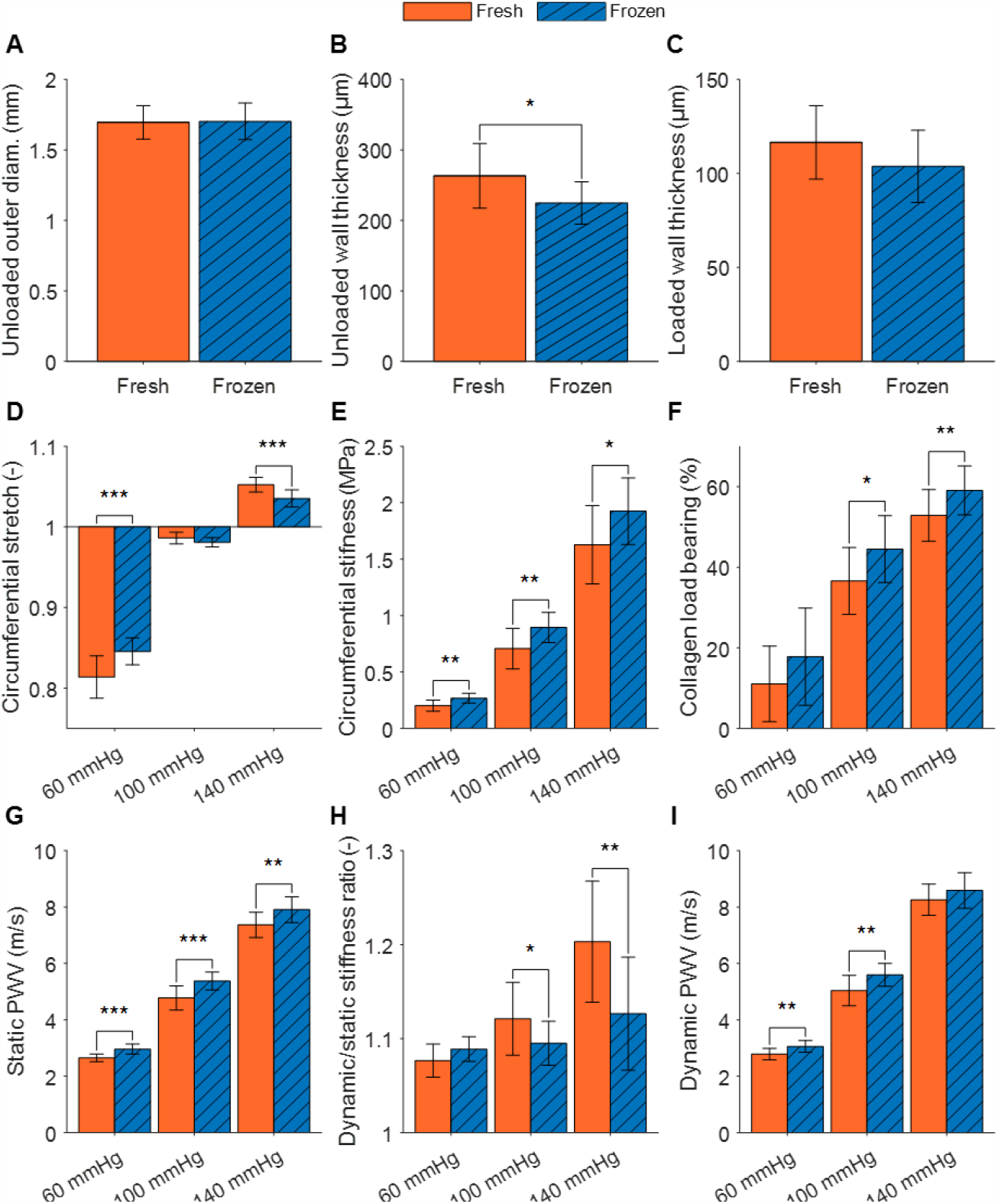
Despite a loss of volume (A, B and C), frozen samples display higher circumferential stiffness than fresh samples (D, E and F,), however less so under dynamic pressurization conditions, when such difference is attenuated by a reduction in viscoelastic stiffening in frozen samples (G, H and I). Panels A and B display results for unpressurized samples at unloaded length, while panel C displays results at 100 mmHg of pressure displays. Panels C through I display results for samples at in vivo-like length, while panels H and I display results for dynamic pressurization conditions using a 10 Hz pulse frequency. * p<0.05, ** p<0.01 and *** p<0.001.

### 3.3 Effect of flash freezing on the dynamic response to inflation

Flash freezing reduced viscoelastic stiffening of the aortic wall during dynamic pressurization (**Figure 4H**). Compared to fresh samples, dynamic-to-quasi-static circumferential stiffness ratios were significantly decreased in flash frozen samples at normotensive and hypertensive pressure ranges. The reduced viscoelastic stiffening meant that the differences in PWV between fresh and frozen samples under dynamic loading conditions were less prominent (**Figure 4I**) compared to those under static conditions (**Figure 4G**), showing no difference between fresh and frozen samples in dynamic PWV at high pressures.

### 3.4 Effect of flash freezing on collagen microstructure

TPSLM image stacks could be obtained up to 80 μm deep into the tissue before losing all contrast. This imaging depth was sufficient to capture the transition from adventitia to media, characterized by a sharp drop in collagen content, but not sufficient to image completely through the arterial wall. Representative maximum intensity projections of the fresh and frozen groups illustrate that aortas displayed a network of wavy, collagen fibers, with no distinct differences between the two groups (**Figure 5A** and **B**). Similarly, flash freezing did not cause significant changes to collagen layer thickness and collagen voxel volume within image stacks (**Figure 5C** and **D**). Despite this, because of the reduced wall thickness of frozen samples (**Figure 4B** and **C**), their collagen volume fraction was significantly higher than that of fresh samples (**Figure 5E**).

**Figure 5:**
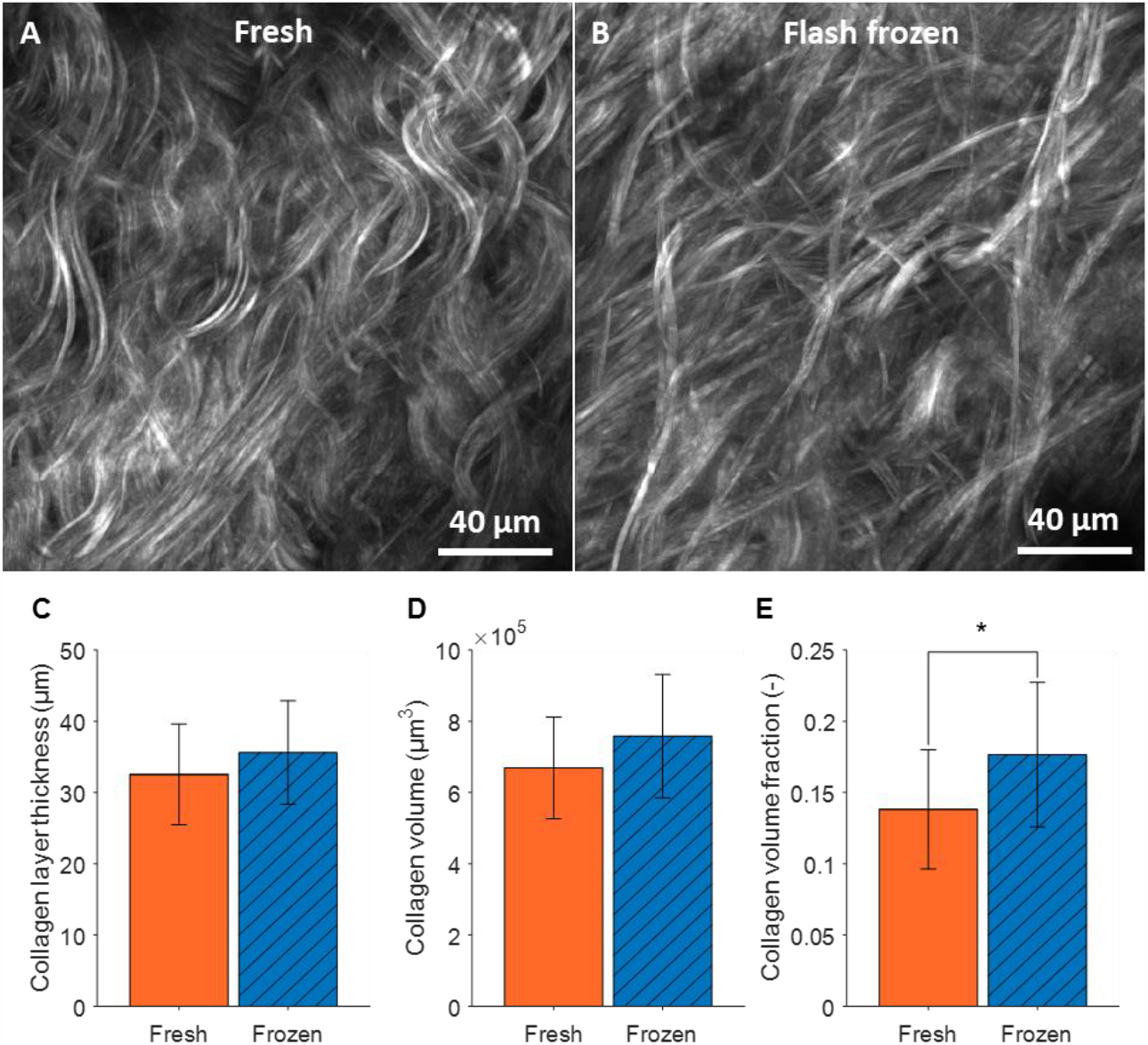
Fresh and frozen samples display similar collagen structures and volume, while frozen samples display an increased collagen volume fraction. Panel A and B display representative maximum intensity projections of second harmonic generation image stacks from fresh and frozen groups under loaded conditions, respectively. Both groups showing similar wavy collagen fibers throughout the arterial wall. Panel C and D show similar collagen layer thickness and volume, respectively, for fresh and flash frozen samples. Panel E shows that frozen samples display a significantly increased collagen volume fraction. * p<0.05.

## 4 DISCUSSION

The present study investigates the effect of flash freezing on the static and dynamic biomechanical properties of the arterial wall tissue. Our results show that flash freezing (1) reduces unloaded arterial wall thickness, (2) increases static circumferential stiffness, (3) increases collagen load bearing, (4) reduces viscoelastic stiffening, and (5) leaves adventitial collagen microstructure intact.

From a mechanical standpoint, our results are in line with previous research, demonstrating that cryopreservation, specifically flash freezing, results in increased static circumferential stiffness (**Figure 4**) [17, 26]. Our constituent-based modeling approach indicated increased collagen circumferential load bearing, primarily at higher pressures, as a potential root cause of these changes. Therefore, in apparent disagreement with our microscopy results (**Figure 5**), increased collagen load bearing may suggest that flash freezing did alter the collagen network in some manner. Previous studies have postulated that thermal cross-linking of collagen fibrils, which alters the collagen network on a sub-fiber scale, might cause circumferential stiffening when flash freezing arterial tissue or constructs [26, 27]. Increased cross-linking would stiffen the collagen network, thereby explaining the increased collagen load bearing, circumferential stiffness and PWV. In contrast, our study shows clear microstructural alterations at adventitial-fiber network level.

Besides the detectable effect of flash freezing on static elastic properties, our results show that flash freezing also alters the viscoelastic properties of the aortic wall. As illustrated in **Figure 4H**, flash frozen samples displayed significantly decreased viscoelastic stiffening at normotensive and hypertensive pressure ranges compared to fresh samples, whilst displaying similar levels of viscoelastic stiffening at hypotensive pressure ranges. Because of the potential effect of cryopreservation on cell integrity and because VSMCs contribute significantly to passive stress relaxation in the arterial wall[28], flash freezing was, indeed, expected to impact viscoelastic stiffening. However, the unchanged viscoelastic properties at low pressures may contradict this hypothesis. Previous work has shown that the VSMCs’ contribution to the passive stress relaxation of arterial wall samples is significant at all levels of applied load [28]. Considering this observation, it might be possible that flash freezing preserve VSMC passive viscoelastic properties to some extent, despite possible dehydration and loss of viability (not studied).

It is interesting to note that, collagen cross-linking may play a key role in reducing viscoelastic stiffening. Increased collagen cross-linking has been shown to reduce intrafibrillar dissipation, the dominant mode of viscous dissipation for collagen-dominated structures, in tissues and collagen constructs [29-31]. Furthermore, collagen has been previously shown to contribute to stress relaxation of the arterial wall, but only after a sufficiently large initial stress is applied [28]. Increased collagen cross-linking could explain the observed flash freezing-induced changes to both static and dynamic passive arterial biomechanics: namely, increased collagen load bearing and circumferential stiffness, as well as reduced circumferential stretch and viscoelastic stiffening at higher pressures.

**Figure 4B** shows that flash freezing reduced the unloaded wall thickness of abdominal aortic segments. Because SHG image stack analysis showed no significant decrease in collagen layer thickness between fresh and flash frozen samples, the loss of wall thickness appeared to be in the media and intima. Freezing-induced cell dehydration or rupture could explain this localized loss of wall thickness, considering that the media and intima are more densely populated by cells than the adventitia [32]. Intima-media thinning rather than increased collagen content appeared to cause increased collagen volume fraction, given similar collagen layer thicknesses for fresh and flash frozen samples. Furthermore, qualitatively, fresh and flash frozen samples displayed similar collagen-network structures, with both groups showing long, wavy, interwoven fibers. Hence, our microscopy findings support the notion that flash freezing did not significantly alter the collagen network organization in the arterial wall.

### 4.1 Limitations

Failure to contain luminal pressure caused uneven sample dropout among the different sample subgroups, resulting in an unbalanced dataset. Because a type III three-way ANOVA analysis was used, our unbalanced datasets resulted in a slight decrease in overall statistical power, while preserving equal weighting among subgroups. Hence, our statistical comparison between fresh and frozen groups remains valid, despite sample dropout.

Considering that our results suggest arterial wall constituents were impacted differently by flash freezing, the use of a spontaneous hypertensive animal model might have skewed our findings. Hypertension is known to trigger remodeling of the arterial wall, leading to increased collagen content, as well as VSMC phenotype switching and contractile dysfunction [33, 34]. Compared to arteries from normotensive animals, possible increased collagen content in arteries of our hypertensive animals could have exacerbated the impact flash freezing.

The potential impact of storage duration on arterial elastic properties was not considered during this study, because not enough tissue was available to measure at multiple storage durations. While literature shows storage duration to affect the total and soluble collagen content of normally frozen (not cryopreserved) arterial tissue, it did not establish a relation between storage duration and altered arterial elastic properties [17]. Like our findings, this study found freezing to have a stiffening effect on arterial tissue, regardless of storage duration. Thus, the freezing process appears to have a larger impact on arterial biomechanics than storage duration.

### 4.2 Recommendations

Gaining an understanding of the alterations caused by flash freezing in arterial tissue is crucial for determining if it enables the possibility of conducting delayed testing on passive arterial biomechanics. Our findings show that flash freezing appears to preserve the contributions of elastin and VSMCs to static and dynamic passive arterial biomechanics, while it alters the contribution of collagen, appearing stiffer but with diminished viscoelastic stiffening. Considering our findings, we propose that the utilization of flash freezing in conjunction with cryostorage serves as an appropriate preservation technique for conducting delayed assessments on passive arterial biomechanics when immediate testing is impractical or undesirable. However, it is essential to exercise caution in future study designs to account for potential systematic errors, as flash freezing might affect experimental groups in varying ways. Furthermore, we recommend to investigate static as well as dynamic elastic behavior when working with flash frozen tissue, as dynamic testing not only aligns better with physiological conditions but also the decreased viscoelastic stiffening may partially offset the static circumferential stiffening caused by cryopreservation.

## 5 CONCLUSION

Flash freezing largely preserves the static and dynamic mechanical behavior of the rat aortic wall but may induce detectable circumferential stiffening and reduced viscoelastic stiffening at increased transmural pressures. If these effects can be considered systematic errors, then flash freezing and cryogenic storage do provide a suitable sample preservation technique for studies investigating arterial biomechanics whenever fresh tissue analyses are not possible or too cumbersome.

## 6 AUTHOR CONTRIBUTIONS

KWFvL: Conceptualization, methodology, investigation, data curation, visualization, formal analysis, writing - original draft, writing - review & editing. KDR: Conceptualization, supervision, project administration, writing - review & editing. SL: Investigation. NJJEB: Methodology, investigation. LvT: investigation. SF: Resources, funding acquisition. TD: Conceptualization, writing - review & editing. BS: Conceptualization, funding acquisition, project administration, supervision, writing - review & editing. AG: Methodology, data curation, formal analysis, software, supervision, writing - review & editing.

