## Supplementary material for "Evaluating flash freezing for preservation of rat abdominal aorta for delayed biomechanical characterization"

Koen W.F. van der Laan, Koen D. Reesink, Sara Lambrichts, Nicole J.J.E. Bitsch, Laura van der Taelen, Sébastien Foulquier, Tammo Delhaas, Bart Spronck, Alessandro Giudici

### Constitutive modeling framework

The arterial wall was assumed to be a thin-walled cylinder, and its viscoelastic mechanical behavior was modeled using a constituent-based quasi-linear viscoelastic modelling framework [1]. Briefly, the elastic response of the wall tissue was modeled as the superimposed contribution of an isotropic elastic matrix reinforced by four families of collagen fibers.

$$\Psi = \Psi_\mathrm{e} + \Psi_\mathrm{c} = \mu\left(I_{1,\mathrm{e}} - 3\right)^{1+\beta} + \sum_{i=1}^{4} \frac{k_1^i}{4k_2^i}\left[e^{k_2^i\left(I_{4,\mathrm{c}}^i - 1\right)^2} - 1\right], \tag{S1}$$

where $\Psi_\mathrm{e}$ and $\Psi_\mathrm{c}$ are the elastin and collagen parts of the wall strain energy density function ($\Psi$), $\mu$ is an elastin stiffness-like parameter, $I_{1,\mathrm{e}}$ is the first invariant of elastin's right Cauchy-Green tensor ($\mathbf{C}_\mathrm{e} = \mathbf{F}^\mathrm{T}\mathbf{G}_\mathrm{e}^\mathrm{T}\mathbf{G}_\mathrm{e}\mathbf{F}$), $\beta$ is a non-neo-Hookean coefficient controlling the functional form of $\Psi_\mathrm{e}$, $k_1^i$ and $k_2^i$ are a stiffness-like and a non-linearity parameter for the *i*-th collagen fiber family, respectively, and $I_{4,\mathrm{c}}^i$ is the fourth invariant of the right Cauchy-Green tensor of the *i*-th collagen fiber family ($\mathbf{C}_\mathrm{c}^i = \mathbf{F}^\mathrm{T}\left(\mathbf{G}_\mathrm{c}^i\right)^\mathbf{T}\mathbf{G}_\mathrm{c}^i\mathbf{F}$). $\mathbf{G}_\mathrm{e}$ is the elastin deposition stretch matrix, which describes elastin's predeformed state in the *in vivo* reference configuration (i.e., at the *in vivo*-like axial stretch and 100 mmHg of pressure)

$$\mathbf{G}_\mathrm{e} = \mathrm{diag}\left[\frac{1}{\lambda_{\theta,\mathrm{e}}\lambda_{z,\mathrm{e}}}, \lambda_{\theta,\mathrm{e}}, \lambda_{z,\mathrm{e}}\right], \tag{S2}$$

where $\lambda_{\theta,\mathrm{e}}$ and $\lambda_{z,\mathrm{e}}$ are elastin's deposition stretches in the circumferential and axial directions, respectively. $\mathbf{G}_\mathrm{c}^i$ is the deposition stretch matrix of the i-th collagen fiber family

$$\mathbf{G}_\mathrm{c}^i = \mathrm{diag}\left[\frac{1}{\lambda_\mathrm{c}^2 \sin\alpha^i \cos\alpha^i}, \lambda_\mathrm{c}\sin\alpha^i, \lambda_\mathrm{c}\cos\alpha^i\right], \tag{S3}$$

where $\lambda_\mathrm{c}$ is the collagen fiber deposition stretch in the fiber direction, and $\alpha^i$ is the *i*-th collagen fiber family's orientation angle in the circumferential-axial plane. $\alpha^1 = 0°$, $\alpha^2 = 90°$, and $\alpha^{3,4} = \pm\alpha$ denote axially, circumferentially, and diagonally oriented fibers respectively.

The viscoelastic behavior of elastin and collagen is governed by two reduced relaxation functions of the type

$$Q_j(t) = \left[1 + \nu_j\left(\int_{\frac{t}{\tau_{2,j}}}^{+\infty} \frac{\mathrm{e}^{-m}}{m}\mathrm{d}m - \int_{\frac{t}{\tau_{1,j}}}^{+\infty} \frac{\mathrm{e}^{-m}}{m}\mathrm{d}m\right)\right]\left[1 + \nu_j \ln\left(\frac{\tau_{2,j}}{\tau_{1,j}}\right)\right]^{-1}, \tag{S4}$$

where $j = \{\mathrm{e}, \mathrm{c}\}$ (e = elastin, c = collagen), and $\nu_j$, $\tau_{1,j} = 0.001$, and $\tau_{2,j}$ are the viscous gain and relaxation time constants of constituent *j*. Given Eqs. S1 and S4, the viscoelastic Cauchy stress at any time *t* can be calculated as

$$\begin{aligned}\sigma_{ii}(t) = \Bigg\{S_{ii}(0) + \int_0^t \Bigg[Q_\mathrm{e}(t-s)\frac{\mathrm{d}}{\mathrm{d}s}\left(-\frac{p_\mathrm{e}}{\lambda_i^2(s)} + 2\frac{\partial\Psi_\mathrm{e}(s)}{\partial\lambda_i^2(s)}\right) + \\ + Q_\mathrm{c}(t-s)\frac{\mathrm{d}}{\mathrm{d}s}\left(-\frac{p_\mathrm{c}}{\lambda_i^2(s)} + 2\frac{\partial\Psi_\mathrm{c}(s)}{\partial\lambda_i^2(s)}\right)\Bigg]\mathrm{d}s\Bigg\}\lambda_i^2(t),\end{aligned} \tag{S5}$$

where $i = \{r, \theta, z\}$, $p_\mathrm{e}$ and $p_\mathrm{c}$ are Lagrange multipliers enforcing incompressibility for the elastin- and collagen-borne part of the wall stress, respectively. The 2nd Piola-Kirchhoff stress at $t = 0$ is

$$S_{ii}(0) = \left(-\frac{p_\mathrm{e}}{\lambda_i^2(0)} + 2\frac{\partial \Psi_\mathrm{e}(0)}{\partial \lambda_i^2(0)}\right)\left[1 + \nu_\mathrm{e} \ln\left(\frac{\tau_{2,\mathrm{e}}}{\tau_{1,\mathrm{e}}}\right)\right]^{-1} + \left(-\frac{p_\mathrm{c}}{\lambda_i^2(0)} + 2\frac{\partial \Psi_\mathrm{c}(0)}{\partial \lambda_i^2(0)}\right)\left[1 + \nu_\mathrm{c} \ln\left(\frac{\tau_{2,\mathrm{c}}}{\tau_{1,\mathrm{c}}}\right)\right]^{-1} \quad \text{(S6)}$$

Given Eq. S5, the luminal pressure and transducer axial force can be calculated as

$$P = (\sigma_{\theta\theta} - \sigma_{rr})\frac{h}{r} \quad \text{and} \quad F_\mathrm{T} = \pi(2\sigma_{zz} - \sigma_{\theta\theta} - \sigma_{rr})hr\,, \quad \text{(S7)}$$

where *h* and *r* are the loaded (i.e., axially stretched and pressurized) wall thickness and mid-wall radius of the vessel.

### Biomechanical variables

We used our constituent-based quasi-linear viscoelastic model to determine biomechanical variables of interest at three different pressure levels: 60, 100, and 140 mmHg, corresponding to hypotensive, normotensive, and hypertensive pressure conditions, respectively. Because the mouse aorta was modeled as a thin-walled cylinder, all estimated variables refer to the mid-wall radial coordinate.

Stresses and stiffnesses were then calculated in the current configuration, i.e., in terms of the Cauchy stress and using the small on large theory [2].

The stored elastic energy per unit length was determined as

$$\Delta\Psi = (\Psi|_\mathrm{SBP} - \Psi|_\mathrm{DBP}) \quad \text{(S8)}$$

where $\rho_\mathrm{o}$ and $\rho_\mathrm{i}$ are the inner and outer radii in the *in vivo* reference configuration, and $\Psi|_\mathrm{SBP}$ and $\Psi|_\mathrm{DBP}$ is the strain energy density function calculated at systolic and diastolic pressure, respectively (Eq. S1). Systolic and diastolic pressure were set to 40–80, 80–120, and 120–160 mmHg for the hypotensive, normotensive, and hypertensive ranges, respectively.

Collagen and elastin's circumferential load bearing were determined as

$$\text{Load bearing} = \frac{\sigma_{\theta\theta,j}}{\sigma_{\theta\theta}} \cdot 100\%$$

where $\sigma_{\theta\theta,j}$ is the contribution to the total stress given by constituent $j = \{\mathrm{e}, \mathrm{c}\}$, as shown in Eq. S5. Finally, pulse wave velocity (PWV) was determined from the simulated static and dynamic pressure diameter relationships using a linearized Bramwell-Hill equation

$$\mathrm{PWV} = \sqrt{\frac{(\mathrm{SBP} - \mathrm{DBP})D_\mathrm{DBP}^2}{\rho(D_\mathrm{SBP}^2 - D_\mathrm{DBP}^2)}}\,, \quad \text{(S9)}$$

where $\rho = 1050\ \mathrm{kg/m^3}$ is the blood density, and $D_\mathrm{SBP}$ and $D_\mathrm{DBP}$ are the artery's inner diameter at the chosen systolic and diastolic pressures, respectively.

**Table S1**: Standard quasi-linear viscoelastic model parameters of the n = 30 Rat abdominal aortas tested in this study

| Sample | Storage condition | Body type | Segment | Elastin | | | | | Collagen | | | | | | | | | |
|---|---|---|---|---|---|---|---|---|---|---|---|---|---|---|---|---|---|---|
| | | | | $\mu$ [kPa] | $\lambda_{\theta,\mathrm{e}}$ [-] | $\lambda_{z,\mathrm{e}}$ [-] | $\nu_{\mathrm{e}}$ [-] | $\tau_{2,\mathrm{e}}$ [s] | $k_1^{1-4}$ [kPa] | $k_2^1$ [-] | $k_2^2$ [-] | $k_2^{3,4}$[-] | $\alpha^{3,4}$ [°] | $\lambda_{\mathrm{c}}$ [-] | $\nu_{\mathrm{c}}$ [-] | $\tau_{2,\mathrm{c}}$ [-] | $R^2_{D}$ [-] | $R^2_{QS}$ [-] |
| I A | Fresh | Lean | Proximal | 23.7 | 1.78 | 1.50 | 0.029 | 100 | 9.7 | 1.09 | 1.92 | 3.60 | 42.8 | 1.27 | 0.022 | 100 | 0.961 | 0.990 |
| II A | | | | 27.6 | 1.72 | 1.52 | 0.000 | 100 | 63.0 | 1.46 | 0.46 | 12.91 | 44.7 | 1.09 | 0.061 | 100 | 0.993 | 0.997 |
| III A | | | | 30.0 | 1.70 | 1.46 | 0.000 | 100 | 20.2 | 1.81 | 1.46 | 3.48 | 41.6 | 1.25 | 0.038 | 100 | 0.989 | 0.995 |
| IV A | | | | 26.1 | 1.76 | 1.49 | 0.000 | 100 | 20.8 | 1.28 | 1.34 | 3.20 | 41.4 | 1.24 | 0.042 | 100 | 0.994 | 0.997 |
| V A | | | | 42.3 | 1.67 | 1.43 | 0.020 | 100 | 15.2 | 1.30 | 1.45 | 3.20 | 42.8 | 1.28 | 0.035 | 100 | 0.989 | 0.996 |
| VI B | | | Distal | 33.9 | 1.65 | 1.50 | 0.000 | 100 | 35.9 | 0.12 | 2.07 | 6.48 | 41.1 | 1.17 | 0.052 | 100 | 0.994 | 0.997 |
| VII B | | | | 36.8 | 1.72 | 1.43 | 0.008 | 100 | 24.5 | 1.08 | 1.34 | 2.79 | 42.9 | 1.28 | 0.061 | 100 | 0.992 | 0.996 |
| VIII B | | | | 36.4 | 1.57 | 1.42 | 0.012 | 100 | 1.0 | 1.69 | 1.50 | 2.61 | 41.6 | 1.41 | 0.029 | 100 | 0.991 | 0.996 |
| IX B | | | | 35.8 | 1.58 | 1.51 | 0.000 | 100 | 34.9 | 2.03 | 4.85 | 11.73 | 42.2 | 1.14 | 0.032 | 100 | 0.992 | 0.996 |
| X A | | Obese | Proximal | 26.2 | 1.65 | 1.59 | 0.000 | 100 | 37.3 | 0.00 | 2.96 | 8.54 | 43.2 | 1.16 | 0.033 | 100 | 0.996 | 0.997 |
| XI A | | | | 48.0 | 1.72 | 1.35 | 0.019 | 100 | 1.0 | 0.26 | 0.32 | 0.47 | 43.0 | 1.80 | 0.028 | 100 | 0.996 | 0.998 |
| XII A | | | | 40.6 | 1.75 | 1.48 | 0.000 | 100 | 42.7 | 2.49 | 2.99 | 7.24 | 42.6 | 1.17 | 0.032 | 100 | 0.991 | 0.996 |
| XIII A | | | | 25.4 | 1.64 | 1.43 | 0.000 | 100 | 38.8 | 1.39 | 5.16 | 11.85 | 44.4 | 1.14 | 0.037 | 100 | 0.989 | 0.997 |
| XIV A | | | | 38.9 | 1.71 | 1.49 | 0.000 | 100 | 37.1 | 2.04 | 2.37 | 5.77 | 41.3 | 1.18 | 0.030 | 100 | 0.994 | 0.997 |
| XV B | | | Distal | 43.3 | 1.75 | 1.35 | 0.036 | 100 | 6.8 | 0.46 | 0.86 | 1.31 | 42.0 | 1.43 | 0.020 | 100 | 0.995 | 0.997 |
| XVI B | | | | 25.1 | 1.71 | 1.52 | 0.000 | 100 | 64.6 | 1.73 | 3.35 | 15.95 | 45.6 | 1.12 | 0.045 | 100 | 0.994 | 0.997 |
| IX A | Frozen | Lean | Proximal | 27.4 | 1.74 | 1.54 | 0.037 | 100 | 20.9 | 0.51 | 0.43 | 1.45 | 45.0 | 1.36 | 0.019 | 100 | 0.983 | 0.994 |
| VI A | | | | 18.8 | 1.67 | 1.65 | 0.000 | 100 | 29.5 | 1.16 | 1.70 | 4.49 | 44.0 | 1.23 | 0.028 | 100 | 0.988 | 0.996 |
| VIII A | | | | 41.3 | 1.64 | 1.40 | 0.018 | 100 | 11.7 | 1.45 | 1.55 | 3.11 | 41.9 | 1.29 | 0.033 | 100 | 0.974 | 0.992 |
| VII A | | | | 31.3 | 1.54 | 1.43 | 0.011 | 100 | 2.2 | 0.53 | 0.62 | 1.07 | 43.3 | 1.53 | 0.040 | 100 | 0.992 | 0.997 |
| I B | | | Distal | 29.2 | 1.59 | 1.46 | 0.000 | 100 | 26.9 | 4.54 | 3.92 | 10.24 | 38.6 | 1.15 | 0.079 | 100 | 0.993 | 0.997 |
| IV B | | | | 19.3 | 1.71 | 1.66 | 0.000 | 100 | 38.9 | 0.61 | 2.60 | 3.85 | 45.8 | 1.20 | 0.061 | 100 | 0.987 | 0.993 |
| V B | | | | 27.0 | 1.63 | 1.48 | 0.010 | 100 | 9.9 | 1.22 | 1.62 | 3.50 | 41.7 | 1.27 | 0.040 | 100 | 0.986 | 0.991 |
| II B | | | | 54.2 | 1.53 | 1.39 | 0.030 | 100 | 14.8 | 1.86 | 3.12 | 5.67 | 44.2 | 1.23 | 0.031 | 100 | 0.982 | 0.994 |
| III B | | | | 40.9 | 1.60 | 1.40 | 0.015 | 100 | 7.0 | 1.46 | 1.82 | 3.43 | 40.9 | 1.30 | 0.030 | 100 | 0.992 | 0.996 |
| XV A | | Obese | Proximal | 18.8 | 1.64 | 1.51 | 0.000 | 100 | 24.5 | 1.67 | 3.13 | 7.28 | 44.4 | 1.18 | 0.019 | 100 | 0.992 | 0.995 |
| XVI A | | | | 38.1 | 1.51 | 1.43 | 0.044 | 100 | 14.0 | 1.83 | 3.99 | 7.95 | 45.7 | 1.20 | 0.025 | 100 | 0.991 | 0.994 |
| X B | | | Distal | 34.0 | 1.63 | 1.44 | 0.000 | 100 | 37.1 | 0.63 | 3.35 | 5.92 | 44.5 | 1.20 | 0.039 | 100 | 0.979 | 0.992 |
| XIII B | | | | 31.0 | 1.58 | 1.46 | 0.000 | 100 | 26.3 | 0.55 | 2.99 | 5.55 | 42.4 | 1.21 | 0.034 | 100 | 0.993 | 0.996 |
| XIV B | | | | 47.4 | 1.59 | 1.43 | 0.053 | 100 | 9.9 | 0.48 | 1.58 | 2.19 | 41.8 | 1.33 | 0.002 | 100 | 0.991 | 0.994 |

$\mu$: elastin stiffness-like parameter, $\lambda_{\theta,\mathrm{e}}$: elastin circumferential deposition stretch, $\lambda_{z,\mathrm{e}}$: elastin axial deposition stretch, $\nu_{\mathrm{e}}$: elastin viscous gain, $\tau_{2,\mathrm{e}}$: elastin time constant 2, $k_1^{1-4}$: collagen fibre stiffness-like parameter, $k_2^1$: collagen non-linearity parameter of the axially oriented fibre family, $k_2^2$: collagen non-linearity parameter of the circumferentially oriented fibre family, $k_2^{3,4}$: collagen non-linearity parameter of the diagonally oriented fibre families, $\alpha^{3,4}$: diagonal collagen fibre orientation angle, $\lambda_{\mathrm{c}}$: collagen deposition stretch, $\nu_{\mathrm{c}}$: collagen viscous gain, $\tau_{2,\mathrm{c}}$: collagen time constant 2, $\mathrm{RSME_D}$: root mean square error of dynamic fitting and $\mathrm{RMSE_{QS}}$: root mean square error of quasi-static fitting.

**Table S2**: Equal-weighted mean model parameters for fresh and frozen sample groups.

| Parameter | Unit | Pressure (mmHg) | Fresh | Frozen | *p*-value |
|---|---|---|---|---|---|
| Unloaded outer diameter | mm | 0 | 1.69 ± 0.12 | 1.70 ± 0.13 | 0.884 |
| Unloaded inner diameter | | | 1.17 ± 0.14 | 1.25 ± 0.13 | 0.132 |
| Unloaded wall thickness | µm | | 263.20 ± 45.93 | 224.72 ± 30.13 | **0.032** |
| Loaded axial stretch | - | 100 | 1.64 ± 0.07 | 1.65 ± 0.07 | 0.848 |
| Loaded outer diameter | mm | | 2.11 ± 0.09 | 2.08 ± 0.08 | 0.388 |
| Loaded inner diameter | | | 1.88 ± 0.08 | 1.88 ± 0.09 | 0.990 |
| Loaded wall thickness | µm | | 116.50 ± 19.46 | 103.74 ± 19.21 | 0.120 |
| Axial stress | MPa | 60 | 0.12 ± 0.02 | 0.12 ± 0.03 | 0.420 |
| | | 100 | 0.15 ± 0.03 | 0.16 ± 0.04 | 0.415 |
| | | 140 | 0.18 ± 0.03 | 0.19 ± 0.05 | 0.439 |
| Axial stiffness | MPa | 60 | 0.58 ± 0.12 | 0.65 ± 0.17 | 0.248 |
| | | 100 | 0.96 ± 0.17 | 1.08 ± 0.23 | 0.145 |
| | | 140 | 1.47 ± 0.23 | 1.60 ± 0.33 | 0.218 |
| Circumferential stretch | - | 60 | 0.81 ± 0.03 | 0.85 ± 0.02 | **0.001** |
| | | 100 | 0.99 ± 0.01 | 0.98 ± 0.01 | 0.087 |
| | | 140 | 1.05 ± 0.01 | 1.04 ± 0.01 | **0.000** |
| Circumferential stress | MPa | 60 | 0.04 ± 0.01 | 0.05 ± 0.01 | **0.039** |
| | | 100 | 0.11 ± 0.02 | 0.12 ± 0.03 | 0.214 |
| | | 140 | 0.18 ± 0.03 | 0.19 ± 0.04 | 0.314 |
| Circumferential stiffness | MPa | 60 | 0.20 ± 0.05 | 0.27 ± 0.04 | **0.003** |
| | | 100 | 0.71 ± 0.18 | 0.89 ± 0.13 | **0.009** |
| | | 140 | 1.63 ± 0.35 | 1.93 ± 0.29 | **0.037** |
| Stored elastic energy | kPa | 60 | 0.71 ± 0.14 | 0.72 ± 0.19 | 0.941 |
| | | 100 | 0.57 ± 0.09 | 0.52 ± 0.16 | 0.265 |
| | | 140 | 0.39 ± 0.07 | 0.38 ± 0.12 | 0.766 |
| Collagen circ. load bearing | % | 60 | 11.05 ± 9.38 | 17.82 ± 12.07 | 0.052 |
| | | 100 | 36.60 ± 8.29 | 44.50 ± 8.27 | **0.006** |
| | | 140 | 52.85 ± 6.39 | 59.03 ± 6.07 | **0.005** |
| D-to-QS stiffness ratio (f=10 Hz) | - | 60 | 1.08 ± 0.02 | 1.09 ± 0.01 | 0.075 |
| | | 100 | 1.12 ± 0.04 | 1.10 ± 0.02 | 0.060 |
| | | 140 | 1.20 ± 0.06 | 1.13 ± 0.06 | **0.005** |
| PWV (static) | m/s | 60 | 2.65 ± 0.13 | 2.96 ± 0.18 | **0.000** |
| | | 100 | 4.77 ± 0.43 | 5.37 ± 0.32 | **0.000** |
| | | 140 | 7.37 ± 0.45 | 7.91 ± 0.46 | **0.008** |
| PWV (f = 10 Hz) | m/s | 60 | 2.79 ± 0.20 | 3.06 ± 0.21 | **0.007** |
| | | 100 | 5.04 ± 0.54 | 5.60 ± 0.40 | **0.006** |
| | | 140 | 8.26 ± 0.56 | 8.59 ± 0.63 | 0.185 |

Data are presented as equal weighted mean ± standard deviation, determined with respect to equal weighted mean. Unloaded parameters were determined at unloaded length while other parameters were determined at in vivo-like length. D-to-QS stiffness ratio: dynamic-to-quasi-static stiffness ratio, circ.: circumferential.

**Table S3**: Equal-weighted mean model parameters for lean and obese sample groups.

| Parameter | Unit | Pressure (mmHg) | Lean | Obese | *p*-value |
|---|---|---|---|---|---|
| Unloaded outer diameter | mm | 0 | 1.71 ± 0.15 | 1.69 ± 0.07 | 0.606 |
| Unloaded inner diameter | mm | 0 | 1.23 ± 0.16 | 1.19 ± 0.10 | 0.420 |
| Unloaded wall thickness | µm | 0 | 239.27 ± 42.53 | 248.65 ± 45.52 | 0.581 |
| Loaded axial stretch | - | 100 | 1.70 ± 0.06 | 1.60 ± 0.05 | **0.000** |
| Loaded outer diameter | mm | 100 | 2.10 ± 0.08 | 2.10 ± 0.09 | 0.957 |
| Loaded inner diameter | mm | 100 | 1.88 ± 0.08 | 1.87 ± 0.09 | 0.589 |
| Loaded wall thickness | µm | 100 | 106.02 ± 20.53 | 114.22 ± 19.29 | 0.310 |
| Axial stress | MPa | 60 | 0.13 ± 0.03 | 0.11 ± 0.02 | **0.034** |
| | | 100 | 0.17 ± 0.04 | 0.14 ± 0.02 | **0.048** |
| | | 140 | 0.20 ± 0.04 | 0.17 ± 0.03 | **0.035** |
| Axial stiffness | MPa | 60 | 0.68 ± 0.16 | 0.55 ± 0.09 | **0.037** |
| | | 100 | 1.09 ± 0.23 | 0.94 ± 0.14 | 0.067 |
| | | 140 | 1.66 ± 0.30 | 1.41 ± 0.19 | **0.026** |
| Circumferential stretch | - | 60 | 0.82 ± 0.03 | 0.83 ± 0.03 | 0.200 |
| | | 100 | 0.98 ± 0.01 | 0.98 ± 0.01 | 0.766 |
| | | 140 | 1.05 ± 0.01 | 1.04 ± 0.01 | 0.095 |
| Circumferential stress | MPa | 60 | 0.05 ± 0.01 | 0.05 ± 0.01 | 0.478 |
| | | 100 | 0.12 ± 0.02 | 0.11 ± 0.02 | 0.259 |
| | | 140 | 0.19 ± 0.04 | 0.17 ± 0.03 | 0.199 |
| Circumferential stiffness | MPa | 60 | 0.24 ± 0.06 | 0.23 ± 0.06 | 0.386 |
| | | 100 | 0.80 ± 0.18 | 0.80 ± 0.20 | 0.905 |
| | | 140 | 1.79 ± 0.35 | 1.76 ± 0.38 | 0.817 |
| Stored elastic energy | kPa | 60 | 0.75 ± 0.18 | 0.67 ± 0.12 | 0.258 |
| | | 100 | 0.60 ± 0.14 | 0.49 ± 0.09 | **0.031** |
| | | 140 | 0.43 ± 0.11 | 0.34 ± 0.06 | **0.049** |
| Collagen circ. load bearing | % | 60 | 13.17 ± 13.61 | 15.70 ± 6.84 | 0.450 |
| | | 100 | 40.02 ± 10.66 | 41.07 ± 7.10 | 0.693 |
| | | 140 | 55.62 ± 8.04 | 56.26 ± 5.55 | 0.751 |
| D-to-QS stiffness ratio (f=10 Hz) | - | 60 | 1.08 ± 0.02 | 1.09 ± 0.01 | 0.438 |
| | | 100 | 1.10 ± 0.03 | 1.12 ± 0.04 | 0.162 |
| | | 140 | 1.17 ± 0.07 | 1.16 ± 0.09 | 0.817 |
| PWV (static) | m/s | 60 | 2.78 ± 0.24 | 2.82 ± 0.21 | 0.579 |
| | | 100 | 4.94 ± 0.45 | 5.21 ± 0.54 | 0.076 |
| | | 140 | 7.47 ± 0.48 | 7.81 ± 0.56 | 0.077 |
| PWV (f = 10 Hz) | m/s | 60 | 2.89 ± 0.27 | 2.96 ± 0.21 | 0.473 |
| | | 100 | 5.18 ± 0.53 | 5.46 ± 0.57 | 0.134 |
| | | 140 | 8.23 ± 0.56 | 8.62 ± 0.62 | 0.117 |

Data are presented as equal weighted mean ± standard deviation, determined with respect to equal weighted mean. Unloaded parameters were determined at unloaded length while other parameters were determined at in vivo-like length. D-to-QS stiffness ratio: dynamic-to-quasi-static stiffness ratio, circ.: circumferential.

**Table S4**: Equal-weighted mean model parameters for proximal and distal sample groups.

| Parameter | Unit | Pressure (mmHg) | Proximal | Distal | *p*-value |
|---|---|---|---|---|---|
| Unloaded outer diameter | mm | 0 | 1.75 ± 0.14 | 1.65 ± 0.07 | **0.043** |
| Unloaded inner diameter | mm | 0 | 1.27 ± 0.15 | 1.15 ± 0.11 | **0.046** |
| Unloaded wall thickness | µm | 0 | 241.54 ± 44.24 | 246.38 ± 44.95 | 0.775 |
| Loaded axial stretch | - | 100 | 1.65 ± 0.07 | 1.65 ± 0.08 | 0.977 |
| Loaded outer diameter | mm | 100 | 2.14 ± 0.07 | 2.05 ± 0.07 | **0.004** |
| Loaded inner diameter | mm | 100 | 1.92 ± 0.06 | 1.83 ± 0.09 | **0.010** |
| Loaded wall thickness | µm | 100 | 110.73 ± 20.57 | 109.50 ± 20.53 | 0.877 |
| Axial stress | MPa | 60 | 0.12 ± 0.03 | 0.12 ± 0.03 | 0.576 |
| | | 100 | 0.16 ± 0.03 | 0.15 ± 0.04 | 0.609 |
| | | 140 | 0.19 ± 0.04 | 0.18 ± 0.05 | 0.503 |
| Axial stiffness | MPa | 60 | 0.62 ± 0.14 | 0.62 ± 0.16 | 0.975 |
| | | 100 | 1.03 ± 0.21 | 1.01 ± 0.21 | 0.797 |
| | | 140 | 1.60 ± 0.30 | 1.47 ± 0.26 | 0.202 |
| Circumferential stretch | - | 60 | 0.82 ± 0.03 | 0.84 ± 0.02 | **0.035** |
| | | 100 | 0.98 ± 0.01 | 0.98 ± 0.01 | 0.438 |
| | | 140 | 1.05 ± 0.01 | 1.04 ± 0.01 | 0.522 |
| Circumferential stress | MPa | 60 | 0.05 ± 0.01 | 0.05 ± 0.01 | 0.906 |
| | | 100 | 0.12 ± 0.02 | 0.11 ± 0.03 | 0.725 |
| | | 140 | 0.19 ± 0.03 | 0.18 ± 0.04 | 0.653 |
| Circumferential stiffness | MPa | 60 | 0.24 ± 0.06 | 0.23 ± 0.05 | 0.760 |
| | | 100 | 0.79 ± 0.20 | 0.81 ± 0.17 | 0.666 |
| | | 140 | 1.79 ± 0.39 | 1.76 ± 0.33 | 0.848 |
| Stored elastic energy | kPa | 60 | 0.72 ± 0.16 | 0.71 ± 0.17 | 0.889 |
| | | 100 | 0.58 ± 0.11 | 0.51 ± 0.14 | 0.142 |
| | | 140 | 0.40 ± 0.08 | 0.37 ± 0.12 | 0.570 |
| Collagen circ. load bearing | % | 60 | 9.11 ± 8.75 | 19.76 ± 9.98 | **0.004** |
| | | 100 | 36.37 ± 7.68 | 44.72 ± 7.71 | **0.004** |
| | | 140 | 52.81 ± 5.95 | 59.07 ± 5.79 | **0.005** |
| D-to-QS stiffness ratio (f=10 Hz) | - | 60 | 1.08 ± 0.02 | 1.08 ± 0.01 | 0.945 |
| | | 100 | 1.11 ± 0.04 | 1.11 ± 0.03 | 0.905 |
| | | 140 | 1.19 ± 0.07 | 1.14 ± 0.08 | 0.060 |
| PWV (static) | m/s | 60 | 2.79 ± 0.21 | 2.82 ± 0.23 | 0.611 |
| | | 100 | 4.93 ± 0.50 | 5.22 ± 0.39 | 0.052 |
| | | 140 | 7.57 ± 0.51 | 7.71 ± 0.54 | 0.463 |
| PWV (f = 10 Hz) | m/s | 60 | 2.90 ± 0.26 | 2.94 ± 0.22 | 0.653 |
| | | 100 | 5.20 ± 0.61 | 5.44 ± 0.42 | 0.218 |
| | | 140 | 8.46 ± 0.62 | 8.39 ± 0.61 | 0.777 |

Data are presented as equal weighted mean ± standard deviation, determined with respect to equal weighted mean. Unloaded parameters were determined at unloaded length while other parameters were determined at in vivo-like length. D-to-QS stiffness ratio: dynamic-to-quasi-static stiffness ratio, circ.: circumferential.
